# Differential effects of BCG-Russia and BCG-TICE on trained immunity: potential implications for bladder cancer immunotherapy

**DOI:** 10.64898/2026.04.17.719184

**Authors:** Richard W. Nauman, Peter A. Greer, Andrew W. Craig, Tiziana Cotechini, D. Robert Siemens, Charles H. Graham

**Author notes:** **Corresponding author:** Charles H. Graham, Department of Biomedical and Molecular Sciences, 18 Stuart Street, Queen’s University, Kingston, Ontario, Canada K7L 3N6.

## Abstract

In recent years, immunotherapy of patients with higher-risk non-muscle invasive bladder cancer (NMIBC) in North America has relied on the use of the TICE strain of BCG. However, limitations in the supply chain have warranted investigation of the therapeutic benefit of other strains of BCG, such as BCG-Russia. Trained immunity, a form of innate immune memory, is now widely believed to be an important component of the therapeutic benefit of BCG. Therefore, in the present study we compared the effects of BCG-TICE and BCG-Russia on the acquisition of trained immunity and related secondary immune responses. C57BL/6 mice received a single intravenous injection of BCG-Russia or BCG-TICE. Four weeks later, bone marrow was collected for flow cytometric analysis of hematopoietic stem and progenitor cell (HSPC) populations, generation of bone marrow-derived macrophages, functional assessment of trained immunity, and transcriptomic profiling. Compared with BCG-Russia, BCG-TICE elicited stronger levels of trained immunity, characterized by higher production of several proinflammatory cytokines upon secondary activation. BCG promoted the expansion of HSPCs independent of strain. BCG-TICE was linked to upregulation of key inflammation-related genes and enrichment of functionally relevant pathways. The results of this study reveal strain-dependent differences in the ability of BCG to induce innate immune memory and inflammatory pathways that could ultimately determine efficacy of immunotherapy of patients with NMIBC.

## Introduction

Since its isolation in 1921, Bacillus Calmette-Guérin (BCG) has been disseminated globally, leading to the emergence of distinct strains.^1^ The TICE strain, preferred for intravesical therapy,^2^ is currently produced at a single facility in the United States. Prior to 2017, the Connaught strain (Sanofi Pasteur) was available as an alternative to BCG-TICE (OncoTICE^®^), but production has since been discontinued. The BCG-Russia strain produced by Verity Pharmaceuticals is approved as an alternative in Canada, but not in the United States. Ongoing shortages underscore the need for direct comparisons of the efficacy of different BCG strains, and several head-to-head clinical trials have explored this.^3-5^ While major differences in clinical outcomes were not reported between most strains, BCG-TICE was shown to extend recurrence-free survival compared to BCG-Connaught,^3,6^ provided patients received maintenance therapy. This benefit did not extend to progression-free survival.^3^ With BCG-Connaught no longer available to patients, the finding of decreased efficacy compared to BCG-TICE has limited relevance. However, with BCG-Russia replacing Connaught as an approved substitute in some jurisdictions when TICE is unavailable, it is crucial to conduct similar comparisons between BCG-Russia and BCG-TICE. The ongoing EVER trial (NCT05037279) aims to address this question.

While BCG is the standard immunotherapy for patients with higher-risk NMIBC, up to 50% of these patients suffer from early disease recurrence,^7-9^ with some progressing to the more lethal muscle-invasive disease.^8,9^ Thus, to develop better therapeutic approaches it is important to improve our understanding of the mechanism of action of BCG. Only recently has it been recognized that BCG induces long-term reprogramming of innate immune cells, a process known as trained immunity. This *de facto* state of immunological memory develops through epigenetic and metabolic reprogramming following recognition of mycobacterial pathogen-associated molecular patterns (PAMPs) by pattern recognition receptors (PRRs).^10^ Trained immunity is thought to contribute to the immunotherapeutic benefit of BCG ^11^ and studies have revealed that trained immunity in peripheral blood monocytes is associated with prolonged recurrence-free survival.^12,13^ Reprogramming of hematopoietic stem cells and myeloid precursors is a key feature of BCG-induced trained immunity,^14^ and these systemic effects of locally administered BCG may be important for achieving maximal therapeutic efficacy.^15^ Recent reports highlight that induction of trained immunity by different strains of BCG results in phenotypic variations.^16^ Thus, strain-dependent effects on the development of trained immunity may have important implications for the management of NMIBC. However, the BCG strains commonly used in NMIBC immunotherapy have not been compared in terms of their ability to induce trained immunity and the trained phenotype elicited. Thus, in the present study, we used a mouse model to assess differential induction of trained immunity and secondary responses between BCG-TICE and BCG-Russia. Our results indicate that functional reprogramming of macrophages differs between the two strains, with stronger secondary immune responses associated with BCG-TICE

## Materials and Methods

### BCG

BCG-Russia was purchased from Verity Pharmaceuticals (Mississauga, ON, CA) as formulated for intravesical instillation (freeze-dried powder, 40 mg net weight per vial, 1-8 × 10^8^ Colony Forming Units (CFU) per vial). BCG-TICE (Merk; Rahway, NJ, U.S.) was obtained from Investigational Drug Services at Kingston Health Sciences Centre (freeze-dried powder, 50 mg net weight per vial, 1-8 × 10^8^ CFU per vial). BCG was aliquoted under aseptic conditions, with minimal exposure to light, and stored at 4°C. Both BCG strains were supplied in vials containing equivalent quantities as measured in CFUs but differed in the net weight of freeze-dried powder. To control for this and standardize the dose to an average of 5.625 × 10^6^ CFU (total amount injected), BCG was resuspended at a concentration of 10 mg/ml (Russian strain) or 12.5 mg/ml (TICE strain) in sterile PBS immediately before use.

### Mouse Models

C57BL/6 mice were purchased from The Jackson Laboratory (Bar Harbor, ME, U.S.) and housed in a pathogen-free facility with *ad libitum* access to food and water. To induce trained immunity, six- to eight-week-old female or male mice were intravenously injected with 5.625 × 10^6^ CFU in 50 μl phosphate-buffered saline (PBS; ThermoFisher; Waltham, MA, U.S.). After four weeks, mice were euthanized via barbiturate overdose and tissues were collected aseptically. All animal experiments were approved by the Queen’s University Animal Care Committee in accordance with the Canadian Council on Animal Care (CCAC) guidelines.

For *in vivo* rechallenge, mice were injected with 10 μg LPS (*E. coli* O26:B6; MilliporeSigma; Burlington, MA, U.S.) intraperitoneally four weeks after BCG administration. Four hours after the LPS injection, whole blood was collected via cardiac puncture in the presence of EDTA to prevent coagulation. Plasma was recovered from the supernatant of centrifuged whole blood and was stored at −80°C. Cytokine quantification was performed by Eve Technologies (Calgary, AB, CA) using the Mouse Focused 10-Plex Discovery Assay^®^ (MDF10). Cytokines in this panel included GM-CSF, IFN-γ, IL-1β, IL-2, IL-4, IL-6, IL-10, IL-12p70, CCL2, and TNF-α. Analytes with concentrations that fell outside the detectable range using the Eve Technologies assay were alternatively measured by ELISA (Invitrogen/ThermoFisher), or their raw fluorescence intensity (FI) values were reported to assess relative levels.

### Ex Vivo Stimulation Assay

Bone marrow was harvested from femurs and tibiae in RPMI-1640 (ThermoFisher) supplemented with 2.5% heat-inactivated FBS (MilliporeSigma). To generate bone marrow-derived macrophages (BMDMs), 2×10^6^ bone marrow cells were plated in 10-cm dishes with 10 mL of BMDM medium (2 mM L-glutamine, 10% FBS, 20 mM HEPES, 1% non-essential amino acids, 1% essential amino acids, 100 U/mL penicillin, 100 μg/mL streptomycin, 1 mM sodium pyruvate, 0.14% 5N NaOH (all ThermoFisher), in RPMI-1640 medium) supplemented with 20 ng/mL of recombinant murine macrophage colony stimulating factor (M-CSF; PeproTech®, ThermoFisher) (BMDM/M-CSF medium). Cells were left to incubate at 37°C and 5% CO_2_ for six days, with an additional 5 mL of BMDM/M-CSF medium added on days two and four. After macrophage differentiation was complete, each dish was washed of residual medium and non-adherent cells. To facilitate detachment from dishes, BMDMs were incubated with 5 mL of Cellstripper (Corning; Corning, NY, U.S.) for 20 minutes. Cells were resuspended in fresh BMDM/M-CSF medium and replated for stimulation assays.

BMDMs seeded in 96-well plates (10^5^ cells/well) were allowed to adhere overnight (37°C, 5% CO_2_) prior to stimulation. After 15-20 hours, the old medium was discarded and fresh medium (containing 10 ng/ml M-CSF) and 100 ng/ml LPS (O26:B6) or 1 μg/ml Pam3CysSerLys4 (Pam3Cys; EMC Microcollections; Tübingen, BW, Germany) was added. After 24 hours, cell-free supernatants were collected and stored at −80°C and cells were lysed for RNA isolation. Cytokine quantification was performed as previously described (MDF10 or ELISA).

### Flow Cytometry

Bone marrow cells (2×10^6^) in suspension were subjected to red blood cell lysis (Pharm Lyse^™^; BD Biosciences; San Diego, CA, U.S.), then incubated (4°C; 30 min.) with anti-CD16/CD32 (1:250; BD Biosciences) and fixable Aqua viability stain (1:500; Invitrogen) in PBS. Subsequently, cells were washed with PBS supplemented with 2 mM EDTA and 0.75% BSA and stained for lineage markers (4°C; 30 min.). Antibodies in the lineage marker cocktail included: anti-Ter-119 (BioLegend; San Diego, CA, U.S.), anti-CD11b (clone M1/70; BioLegend), anti-IgM (clone RMM-1; BioLegend), anti-CD3 (clone 17A2; BioLegend), anti-CD45R (clone RA3-6B2; BioLegend), and anti-Ly6G/C (Gr-1; clone RB6-8C5; BD Biosciences), all FITC-conjugated and at a 1:400 dilution (0.125 μg). Cells were then washed with PBS/EDTA/BSA and stained for HSPC markers (4°C; 30 min.) with 10% Brilliant Stain Buffer (BD Biosciences). The following fluorophore-conjugated antibodies were used for HSPC staining: anti-c-Kit–BV605 (1:400; clone 2B8; BD Biosciences), anti-Sca-1–PE-Cy7 (1:400; clone D7; BD Biosciences), anti-CD150– BV421 (1:200; clone Q38-480; BD Biosciences), anti-CD48–AF700 (1:200; clone HM48-1; BioLegend), anti-Flt3–PE (1:200; clone A2F10.1; BD Biosciences), and anti-CD34–AF647 (1:200; clone RAM34; BD Biosciences). Cells were subsequently washed in PBS/EDTA/BSA and resuspended in Fixation Buffer (BD Biosciences) containing 4.21% formaldehyde for 15 minutes (4°C). After a final wash, cells were resuspended in PBS/EDTA/BSA and stored at 4°C until they were analysed using a CytoFLEX S flow cytometer (Beckman Coulter; Brea, CA, U.S.). Data were analysed using FlowJo software (v10.10; BD).

### Transcriptomic Analysis

RNA was extracted from BMDMs using the PureLink Micro-to-Midi Total RNA Purification kit (Invitrogen). RNA integrity (all samples RIN = 10) was assessed with the Agilent 2100 Bioanalyzer (Agilent Technologies; Santa Clara, CA, U.S.) and quantified using a Nanodrop 2000 spectrophotometer (ThermoFisher). The nCounter assay was performed at the Ontario Institute for Cancer Research (OICR; Toronto, ON, CA) according to the manufacturer’s instructions, using the NanoString Myeloid Innate Immunity V2 Code Set (Bruker Spatial Biology; Bothell, WA, U.S.) and 50 ng RNA input. Data were analyzed with nSolver software (v4.0; Bruker Spatial Biology) and nCounter Advanced Analysis plugin (v2.0). Default parameters were used for normalization, with automated selection of housekeeping genes to minimize pairwise variation according to the geNorm algorithm. Adjusted p-values were generated using the Benjamini– Hochberg method of false discovery rate estimation. Figures were generated with R (v4.5.1)

### Statistical Analysis

Final data analysis was performed using GraphPad Prism (v10.4). One-way ANOVA followed by Tukey’s multiple comparisons test was used to determine statistical differences between three groups. Two-way ANOVA with Šídák’s *post-hoc* analysis was performed on *ex vivo* rechallenge data. The ROUT method (Q = 1%) was used where specified in figure captions to identify outliers, which were excluded from analysis. Data are presented as mean ± SEM, with *p* < 0.05 considered statistically significant.

## Results

### BCG-TICE and BCG-Russia Induce Expansion of Hematopoietic Stem Cells

A known feature of BCG-induced trained immunity is induction of myelopoiesis, with expansion of HSPCs (LKS^+^ cells) peaking four weeks after BCG administration.^14^ Four weeks after an intravenous injection of BCG, both strains induced a marked expansion of LKS^+^ cells compared to controls, with 13.2-fold and 11.2-fold mean increases following BCG-Russia and BCG-TICE vaccination, respectively (Figure 1A, B). Bone marrow proportions of LKS^+^ cells showed a similar pattern, increasing significantly following administration of either strain (Figure 1C). No significant differences in the induction of myelopoiesis were detected between the two BCG strains.

**Figure 1.**
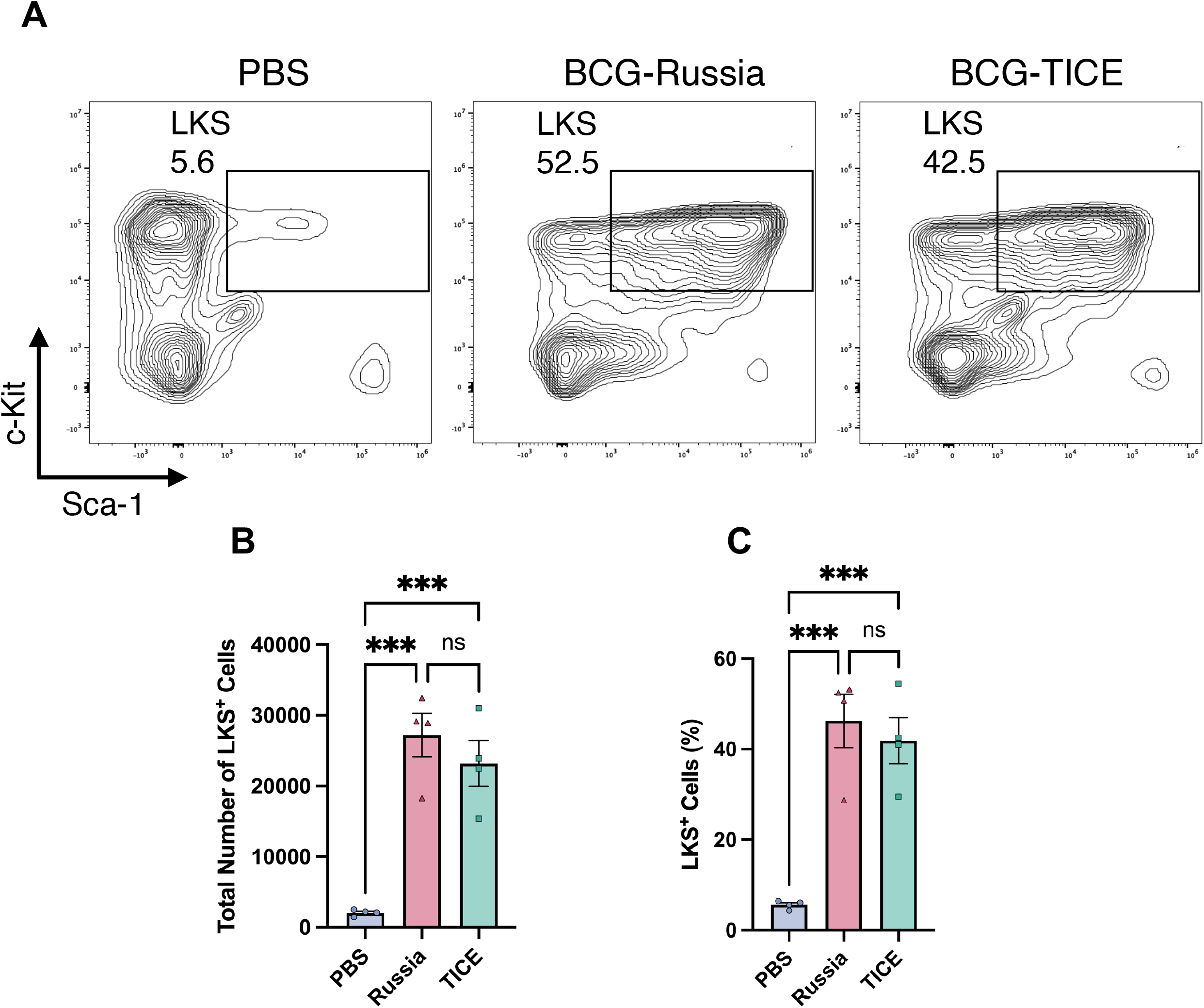
BCG-mediated shifts in HSPC populations. (A) Representative fluorescence-activated cell sorting (FACS) plots of LKS^+^ (Lin^**–**^c-Kit^**+**^Sca-1^**+**^) frequencies four weeks after vaccination. Values on plots indicate LKS^+^ cells as proportion of all Lin^−^ cells. (B) Absolute counts and (C) frequencies of LKS^+^ cells as proportion of all Lin^−^ cells, four weeks after vaccination. Data points represent values from independent biological replicates. Mean ± SEM, one-way ANOVA (ns = not significant, ^***^p < 0.001).

### BCG-TICE is Associated with Elevated Plasma Cytokines Following Secondary In Vivo Challenge with LPS

To assess cytokine responses to heterologous stimulation, mice were challenged intraperitoneally with LPS four weeks after BCG administration. Compared with unvaccinated controls, both BCG-Russia and BCG-TICE vaccination significantly increased IL-1β and IL-12 plasma levels four hours after intraperitoneal challenge with LPS (Figure 2A, B). Mice vaccinated with BCG-TICE exhibited significantly higher TNF-α, IL-1β, and IFN-γ plasma concentrations than mice vaccinated with BCG-Russia (Figure 2C; Supplemental Figure 1). However, there were no significant differences in plasma IL-10 and IL-12 levels in mice vaccinated with BCG-TICE versus BCG-Russia (Figure 2D, E). These findings demonstrate that both BCG strains trigger enhanced cytokine responses to secondary challenge, with BCG-TICE invoking a broader pro-inflammatory profile than BCG-Russia.

**Figure 2.**
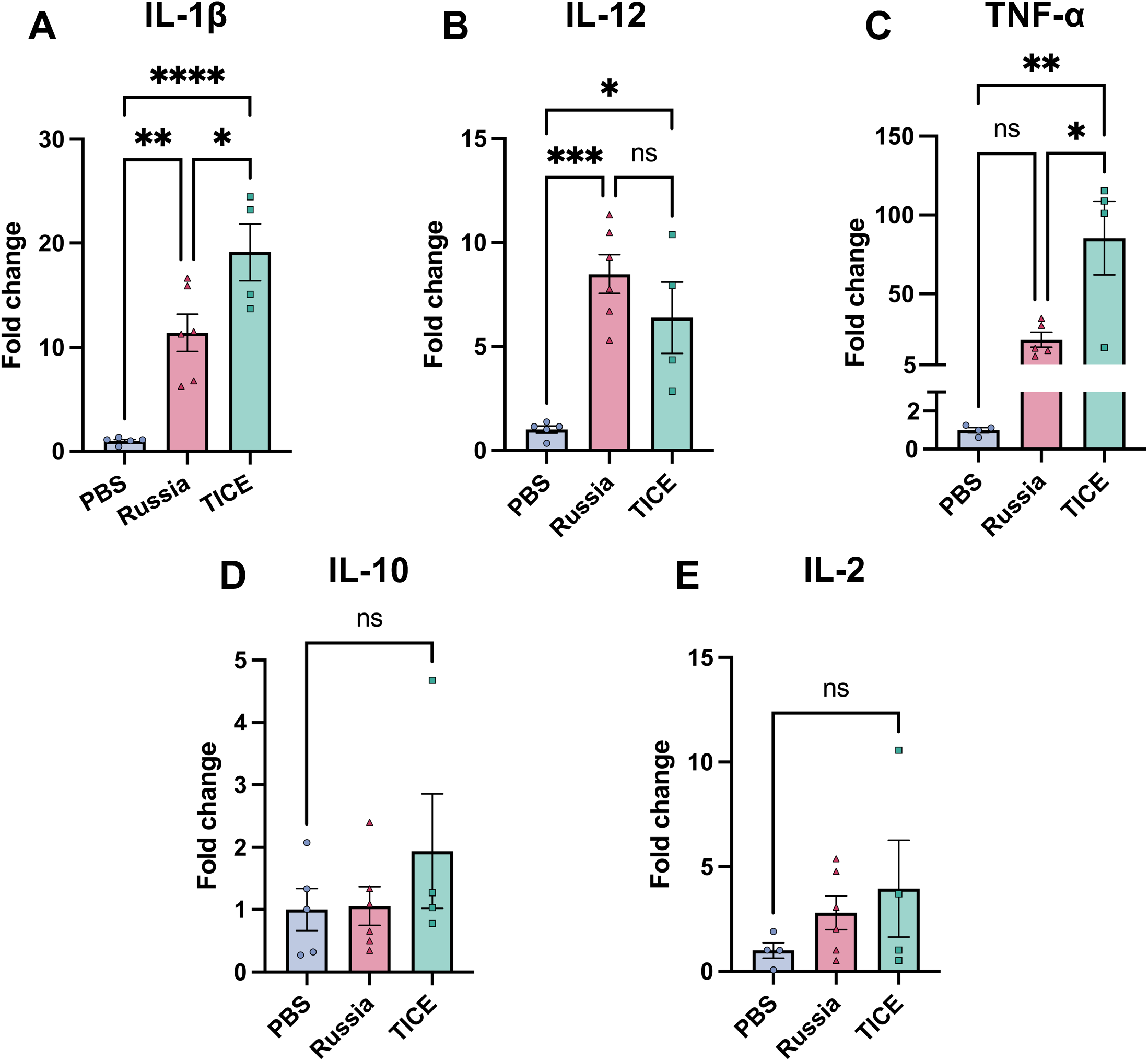
Strain-dependent effects of BCG on plasma cytokine levels. (A-E) Plasma concentrations of IL-1β, IL-12, TNF-α, IL-10, and IL-2 four weeks after PBS or BCG vaccination and four hours after re-challenge (LPS, intraperitoneally). TNF-α concentrations were assessed by ELISA, while all other reported cytokines were measured via the MDF10 multiplex assay. Data points represent values from individual mice (all female). Concentration values from independent experimental repeats were normalized to the control group mean (set to 1.0) and expressed as fold changes. One outlier was removed from the TNF-α data set following an outlier test (ROUT method; Q = 1%). Mean ± SEM, one-way ANOVA (ns = not significant, ^*^p < 0.05, ^**^p < 0.01, ^***^p < 0.001, ^****^p < 0.0001).

### BCG-TICE Elicits Stronger Pro-Inflammatory Cytokine Responses in Macrophages

Acquisition of trained immunity in response to BCG is well-defined in cells of the monocyte/macrophage lineage. Moreover, results from *in vivo* challenge of BCG-trained mice suggested a dominant role of monocytes and macrophages in responding to a secondary stimulus based on the heightened secretion of inflammatory mediators. To further investigate functional reprogramming of macrophages by the two BCG strains, we generated bone marrow-derived macrophages (BMDMs) and measured cytokine production following *ex vivo* restimulation. Compared to BCG-Russia or control groups, BMDMs from BCG-TICE–vaccinated mice exhibited significantly higher production of GM-CSF, IL-1β, IL-6, and TNF-α after a 24-hour stimulation with Pam3Cys (Figure 3A–D). Pam3Cys is a synthetic toll-like receptor (TLR) 1/2 agonist. No differences in CCL2, IL-10, or IL-12 levels were observed across groups after Pam3Cys stimulation (Figure 3E-G). In contrast, a 24-hour stimulation with LPS (a TLR4 agonist) resulted in significantly increased secretion of TNF-α by BMDMs isolated from mice vaccinated with BCG-TICE compared with BMDMs from mice vaccinated with BCG-Russia, whereas the secretion of IL-1β, CCL2, and IL-10 was reduced in cultures of BMDMs from mice vaccinated with BCG-TICE relative to BCG-Russia. IL-12 levels in cultures of BMDMs from mice vaccinated with BCG-TICE were significantly increased compared with BMDMs from unvaccinated mice; however, levels of IL-12 were not significantly different in cultures of BMDMs obtained from mice vaccinated with BCG-TICE relative to BCG-Russia. Collectively, these findings indicate that BCG-TICE induces stronger cytokine responses than BCG-Russia after exposure certain stimuli, but also suppresses specific cytokines, indicating strain-dependent and stimulus-specific training effects.

**Figure 3.**
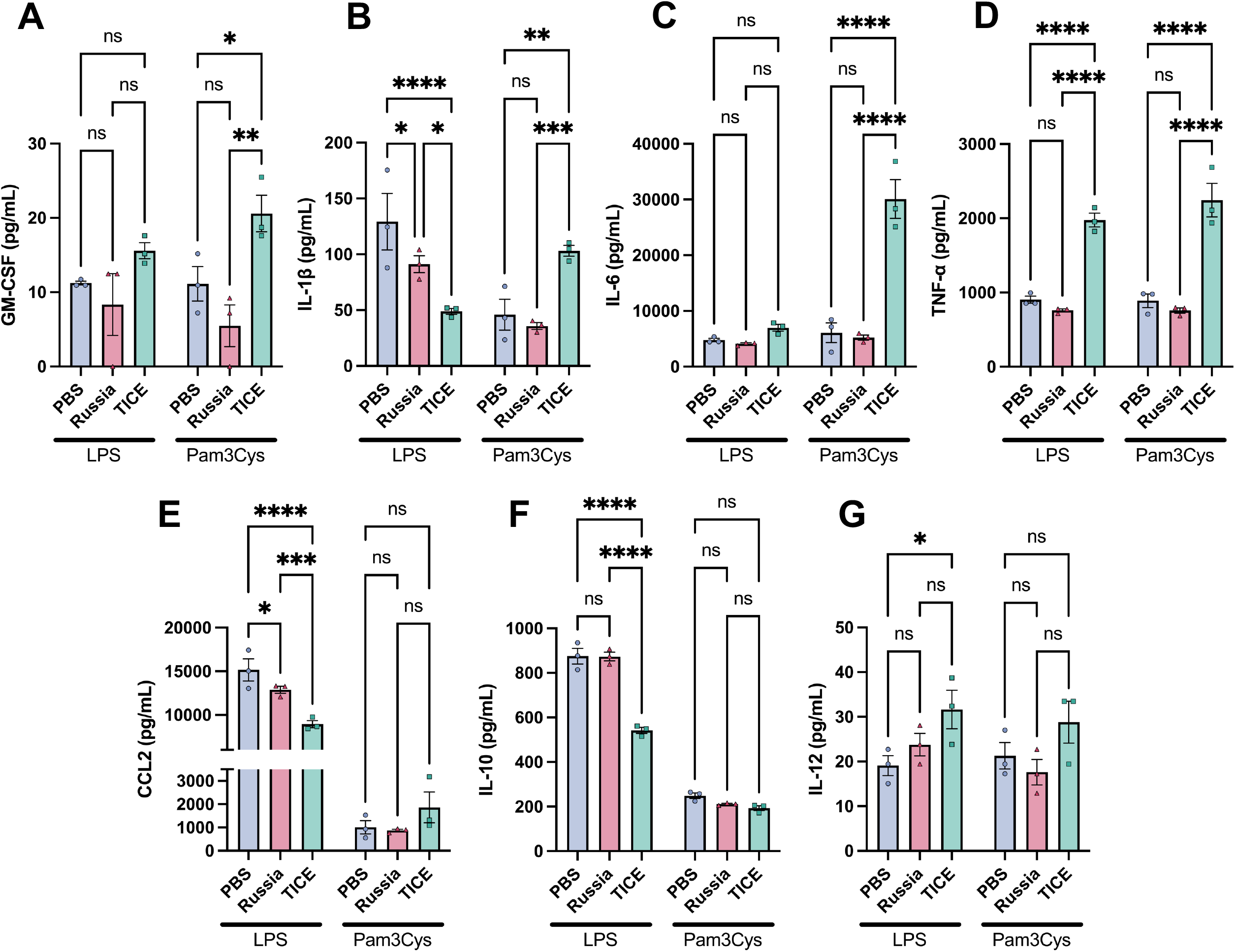
Effects of BCG strain on cytokine production by bone marrow-derived macrophages (BMDMs). (A-G) Concentrations of GM-CSF, IL-1β, IL-6, TNF-α, CCL2, IL-10, and IL-12 in supernatants of BMDMs from PBS- or BCG-vaccinated mice, 24 hours after restimulation. IL-6 concentrations were assessed by ELISA, while all other reported cytokines were measured via the MDF10 multiplex assay. Data points represent values from independent biological replicates (all male). Mean ± SEM, two-way ANOVA (ns = not significant, ^*^p < 0.05, ^**^p < 0.01, ^***^p < 0.001, ^****^p < 0.0001).

### Training with BCG-TICE Triggers Broad Changes in the Expression of Myeloid-related Genes and Pathways Upon Reactivation

Given the divergent effects of the two strains of BCG on the release of inflammatory cytokines, we next determined whether transcriptional profiles in BMDMs were differentially regulated by BCG-TICE and BCG-Russia. We analyzed the expression of a targeted subset of myeloid genes in BMDMs using NanoString nCounter. *In vivo* training with BCG-Russia induced minimal transcriptional changes in LPS-restimulated macrophages, with just two genes (*Ptprc, Gpr65*) differentially expressed compared to controls (Figure 4A). Pathway analysis revealed limited enrichment, with single-gene representation in processes such as cell migration/adhesion and lymphocyte activation (Figure 4B). In contrast, BCG-TICE induced broad transcriptional reprogramming, with 127 genes differentially expressed relative to controls (Figure 4C). The most strongly upregulated genes (*H2-Aa, CD74, H2-Ab1*) were linked to processes such as antigen presentation and cell migration/adhesion. Several other pathways were highly enriched, with Th1 activation, angiogenesis, and cell migration/adhesion having the highest proportions of significantly differentially expressed genes (gene ratio > 0.38; Figure 4D). Notably, every pathway contained at least two differentially expressed genes, with some having over 30 significant genes.

**Figure 4.**
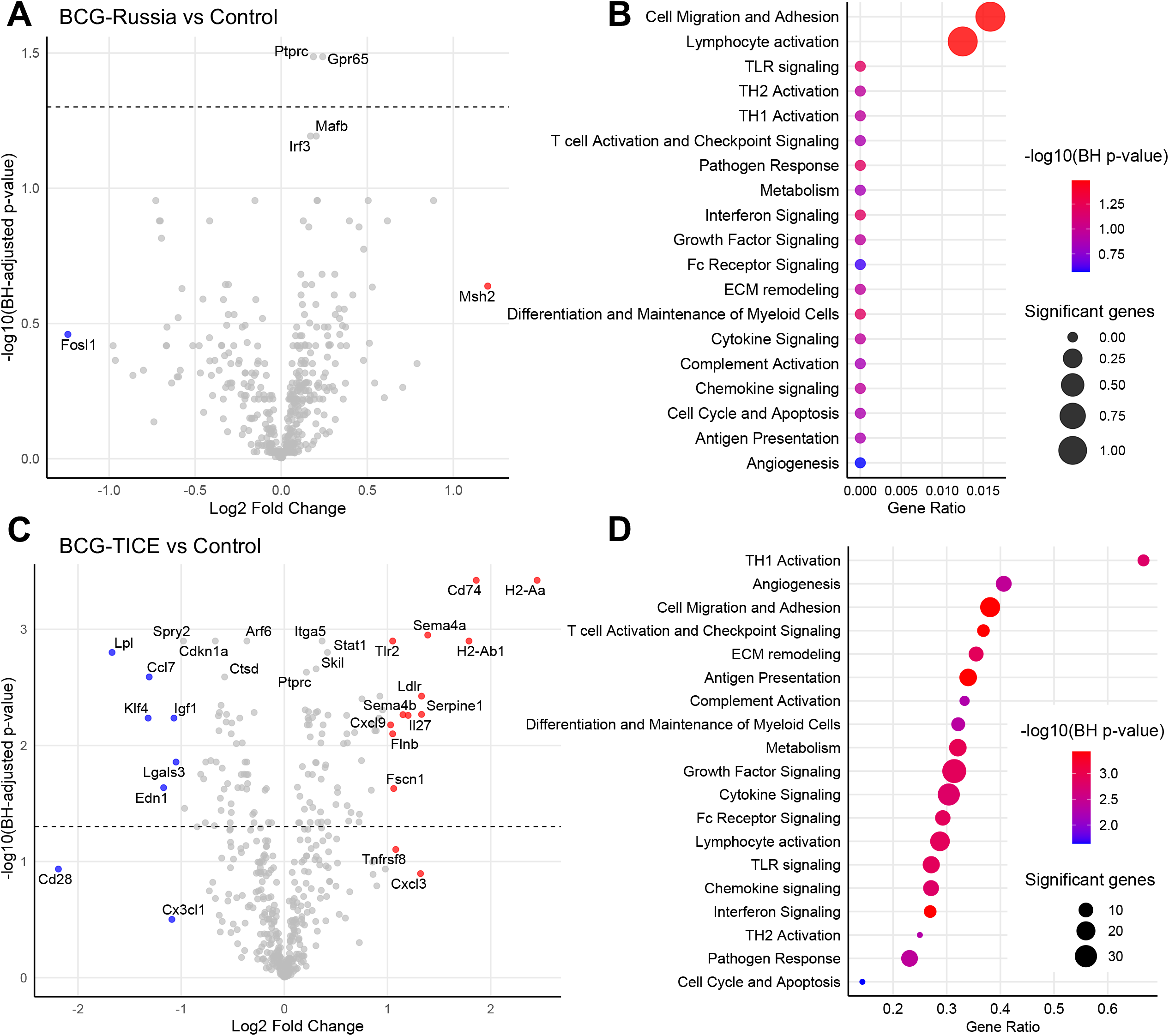
Differentially expressed genes and pathway enrichment in BCG-trained, compared to untrained, macrophages. (A) Volcano plot showing differentially expressed genes and (B) dot plot indicating pathway enrichment in BMDMs from mice trained with BCG-Russia compared to mice administered PBS (untrained controls). (C) Volcano plot showing differentially expressed genes and (D) dot plot indicating pathway enrichment in BMDMs from mice trained with BCG-TICE compared to mice administered PBS. Horizontal dashed line on volcano plots indicates Benjamini– Hochberg (BH) adjusted p-value of 0.05. Genes with log2FC < −1 are shaded blue; genes with log2FC > 1 are shaded red. In dot plots, colour scale is based on the minimum −log10(BH p-value) among genes represented in each pathway. Dot size is proportional to the total number of significant genes in each pathway. Gene ratio on the x-axis accounts for the number of genes that comprise each pathway, expressing the ratio of significant genes as a proportion of all pathway genes. Significant genes may be upregulated or downregulated.

Directed global significance scores (dGSS) were low when comparing BCG-Russia to untrained macrophages, with only interferon signaling, Th1 activation, and complement activation showing modest increases (dGSS 2.1–2.6), none of which contained significantly upregulated genes (Figure 5A). In contrast, 84% of the pathways that emerged in our analysis were upregulated following BCG-TICE vaccination. The highest degree of upregulation was observed in T cell activation and checkpoint signalling (dGSS = 6.2), interferon signalling (dGSS = 5.1), and Th1 activation (dGSS = 4.8), all containing multiple significantly upregulated genes. Notably, several pathways showed opposing regulation, with pathways downregulated with BCG-Russia (*e*.*g*., Th2 activation) showing upregulation with BCG-TICE, whereas pathways suppressed by BCG-TICE (*e*.*g*., cell cycle and apoptosis) showing modest upregulation in BMDMs from mice vaccinated with BCG-Russia.

**Figure 5.**
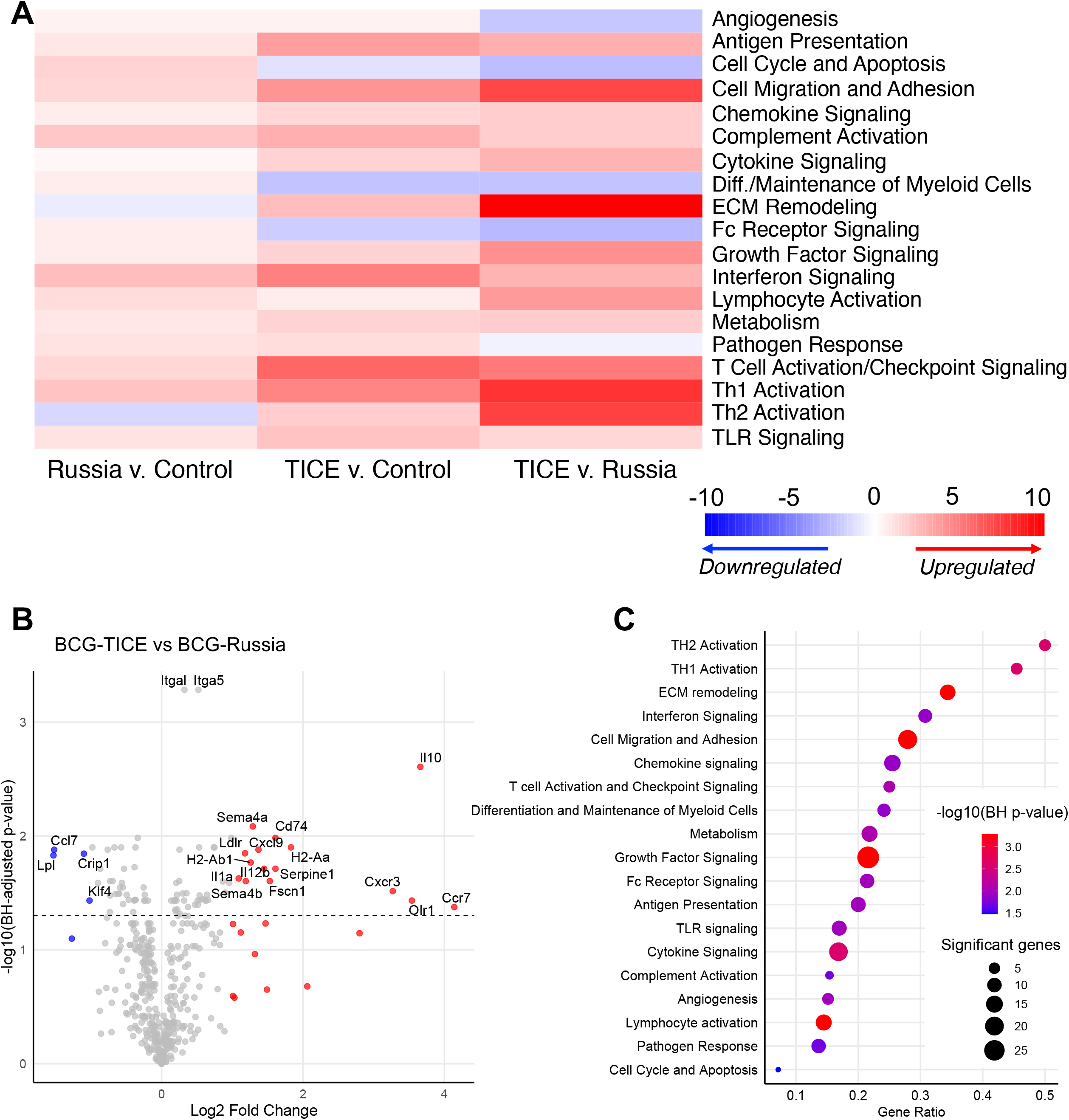
Differentially expressed genes and pathway enrichment between BCG-TICE and BCG-Russia. (A) Heatmap displaying directionality of pathway enrichment for each comparison. Values represent nSolver directed global significance scores. (B) Volcano plot showing differentially expressed genes and (C) dot plot indicating pathway enrichment in BMDMs from mice trained with BCG-TICE compared to mice trained with BCG-Russia. Horizontal dashed line on the volcano plot indicates Benjamini–Hochberg (BH) adjusted p-value of 0.05. Genes with log2FC < −1 are shaded blue; genes with log2FC > 1 are shaded red. In the dot plot, colour scale is based on the minimum −log10(BH p-value) among genes represented in each pathway. Dot size is proportional to the total number of significant genes in each pathway. Gene ratio on the x-axis accounts for the number of genes that comprise each pathway, expressing the ratio of significant genes as a proportion of all pathway genes. Significant genes may be upregulated or downregulated.

Direct comparison of the two BCG strains revealed 51 significantly upregulated genes and 37 significantly downregulated genes in BMDMs from mice vaccinated with BCG-TICE (Figure 5B). Pathways related to Th2 activation, Th1 activation, and ECM remodeling had the greatest representation of significantly differentially expressed genes (Figure 5C). Enrichment in ECM remodeling, cell migration/adhesion, growth factor signalling, and lymphocyte activation was driven by genes with the highest significance. The pathways most strongly upregulated included ECM remodeling (dGSS = 10.4), Th1 (dGSS = 8.3) and Th2 (dGSS = 7.5) activation, and cell migration/adhesion (dGSS = 7.5; Figure 5A). These data indicate that vaccination with BCG-TICE elicits significantly greater transcriptional changes in immune-related genes of macrophages, representing more robust reactivation than vaccination with BCG-Russia.

## Discussion

Divergent immunogenic properties among BCG strains are well characterized and recent studies have extended these differences to trained immunity.^16^ In the present study, we investigated differences in trained immunity phenotypes induced by two strains of BCG used for the treatment of patients with NMIBC. The results revealed clear differences in the abilities of BCG-TICE and BCG-Russia to induce trained immunity, with BCG-TICE eliciting higher levels of trained immunity, demonstrated by enhanced inflammatory responses to subsequent stimuli. Following an *in vivo* heterologous inflammatory challenge, plasma cytokines were most elevated in mice that were vaccinated with BCG-TICE, despite both strains having similar effects on HSPC expansion. This suggests independent mechanisms governing myelopoietic induction versus functional reprogramming during trained immunity acquisition. When macrophages from mice vaccinated with BCG-TICE were re-challenged *ex vivo*, they exhibited markedly enhanced secondary cytokine responses compared to those trained with BCG-Russia. Divergent transcriptional signatures further supported distinct trained immunity profiles between the two BCG strains. Recent work has made progress in establishing the causal relationship between trained immunity and the therapeutic benefit of BCG,^17^ supporting our suggestion that BCG-TICE may be linked to better patient outcomes based on our findings.

Our observation that both BCG-Russia and BCG-TICE trigger an expansion of hematopoietic progenitors in the bone marrow is concordant with previous studies.^14,16^ Despite gross shifts in HSPC populations being a non-specific measure of trained immunity, they are consistent with the systemic effects of intravesical BCG administration triggering myelopoiesis, a critical aspect sustained innate immune memory.^14^ Mature leukocytes that encounter BCG and readily acquire trained immunity may be short-lived, as is the case with circulating monocytes. Bone marrow HSPCs are known to undergo epigenetic and metabolic changes in response to BCG, leading to a long-lasting memory phenotype that is retained during cell division and differentiation. Consequently, trained bone marrow harbors a reservoir of myeloid cells, especially monocytes, that enter the circulation and exert robust inflammatory responses upon reactivation. Our findings reveal that divergence in functional reprogramming, *i*.*e*. trained immunity, following exposure to different strains of BCG can be acquired after myeloid expansion has occurred.

Results of plasma cytokine analysis following *in vivo* LPS re-challenge provided further evidence that the two strains of BCG exert differential effects on functional reprogramming. The pattern of cytokines detected in elevated quantities following BCG-TICE–training is consistent with a monocyte/macrophage driven response. Significant differences between the two strains were seen in the levels of certain acute-phase cytokines (IL-1β, TNF-α, IFN-γ). IL-12 levels were significantly increased in BCG-treated mice compared to controls, but did not differ between the two strains, which could be explained by reduced susceptibility of dendritic cells to strain-dependent effects on trained immunity. Dendritic cells are known to be the main source of plasma IL-12.^18-20^ Moreover, we did not observe increased IL-12 production by trained macrophages that mirrored the heightened plasma concentrations, suggesting a minimal contribution of macrophages to systemic IL-12 concentrations. While it has been suggested that dendritic cells can acquire trained immunity,^15,21^ it remains to be determined whether this was the case in our model.

*Ex vivo* restimulation of BMDMs with LPS or Pam3Cys resulted in divergent cytokine profiles. This may be explained by the distinct PRRs that are selectively targeted by LPS (TLR4) versus Pam3Cys (TLR1/2). Notably, expression of *Tlr2* was significantly upregulated in BCG-TICE– compared to BCG-Russia–trained BMDMs. It is hypothesized that, in the clinical setting, tumour-associated immunogens are responsible for triggering secondary responses in trained myeloid cells.^11^ These molecules may act on a diverse repertoire of PRRs leading to robust reactivation, consistent with what was observed following TLR1/2 stimulation. Our findings were restricted to isolated stimulation of specific TLRs, whereas tumour damage-associated molecular patterns (DAMPs) can act on multiple PRRs concomitantly and chronically. However, TLR4 agonists are appropriate for this single-TLR approach as TLR4 is engaged by several DAMPs (*e*.*g*., HMGB1, heat-shock proteins) that are present in the tumour microenvironment.

In contrast with a similar study by Xu *et al*.,^16^ our analysis revealed more limited transcriptional differences associated with BCG-Russia relative to untrained controls. A key methodological distinction between our study and the study of Xu *et al*. is that we assessed gene expression in re-challenged cells, whereas Xu et al, reported RNA sequencing of BMDMs trained *ex vivo*, without a secondary stimulus. We believe that assessing gene expression after restimulation is a strength of our study, as this reactivated state captures a more relevant trained immunity phenotype, rather than a trained, resting state.

In our study, macrophages from mice administered BCG-Russia displayed remarkably few alterations upon reactivation, with *Ptprc* and *Gpr65* emerging as the only significantly differentially expressed genes, each with a fold-change < 0.25. The most differentially expressed genes when comparing BCG-TICE to control drove enrichment in several important pathways, with at least two of *Sema4a, Cd74, H2-Ab1*, and *H2-Aa* being represented in cell migration/adhesion, Th1 activation, interferon signaling, antigen presentation, and T cell activation and checkpoint signaling. T cells play a central role in the therapeutic mechanism of BCG,^22,23^ with optimal efficacy relying on Th1 immunity.^24,25^ Furthermore, interferon signaling in macrophages promotes polarization to a pro-inflammatory, M1-like, anti-tumoral phenotype and is responsible for directly stimulating cytotoxicity against bladder cancer cells.^26-28^ This broadly supportive impact on anti-tumour immunity may not be limited to comparison against untrained cells, as pronounced upregulation of the Th1 and T cell activation pathways was also observed when directly comparing BCG-TICE to BCG-Russia. The high magnitude of ECM remodelling and cell migration/adhesion upregulation was primarily driven by high statistical significance of *Itgal* and *Itga5* differential expression, each involved in both pathways. *Itgal* and *Itga5* encode the alpha subunit (CD11a) of heterodimeric lymphocyte function-associated antigen 1 (LFA-1) and integrin alpha-5, respectively. Binding of CD11a to intercellular adhesion molecules (ICAMs) is important for macrophage migration to lymph nodes.^29^ It is possible, therefore, that *Itgal* upregulation enhances tumour antigen presentation by pro-inflammatory macrophages in secondary lymphoid tissue. The high-level consequences of ECM remodelling and cell migration by tumour-associated macrophages has been proposed to be a tumour-promoting effect because these cells often induce immune-evasive phenotypes in cancer cells.^30^ However, adequate response to BCG in patients is associated with pro-inflammatory macrophages driving tumour clearance.^30-32^ Taken together with upregulation of processes that promote Th1 immunity and T cell-mediated killing, we posit that enhanced macrophage infiltration in this context enhances anti-tumour immunity.

Overall, our study indicates that optimal induction of trained immunity is favoured by BCG-TICE, compared to BCG-Russia, as evidenced by more robust secondary responses. However, the precise mechanism by which BCG-TICE induces higher levels trained immunity is not known. Other groups have suggested a link to linoleic acid metabolism,^16^ but the genetic variations between strains that translate to stronger or weaker initial reprogramming have yet to be identified.

While we delivered BCG intravenously for strong and reliable induction of trained immunity, this is not a viable approach in the clinical setting due to the high risk of sepsis. This model was favoured for strong, reliable induction of central trained immunity, eliminating the variable reprogramming that results when mice are administered intravesical BCG. Novel approaches to optimizing acquisition of central trained immunity are currently under investigation. Delivery of purified components of the BCG cell wall and nanoparticle formulations are under investigation.^33^ In addition, adjuvants for optimal trained immunity induction from intravesical BCG, such as β-glucan or linoleic acid,^16,34^ have been suggested. While still preserving the important local effects of BCG therapy, these novel therapeutic strategies have the potential to promote consistent and stronger trained immunity acquisition in NMIBC patients.

In conclusion, the present report provides novel evidence of superior trained immunity responses linked to BCG-TICE, compared to BCG-Russia. These findings inform important considerations for treating patients with high-risk NMIBC given the potential role of trained immunity in the mode of action of BCG. Systemic immune activation is critical for effective immunotherapy, including intravesical therapy. Further studies and clinical trials, like the ongoing EVER trial (NCT05037279), are needed to confirm the impact of BCG strain on patient outcomes.

## Author Contributions

*Conceptualization:* RWN and CHG. *Study Design/Methodology:* RWN, TC, and CHG. *Data Acquisition:* RWN. *Data Analysis:* RWN and TC. *Data Interpretation:* RWN and CHG. *Writing – Original Draft:* RWN and CHG. *Writing – Review and Editing:* RWN, PAG, AWC, DRS, TC, and CHG. *Funding Acquisition:* PAG, AWC, DRS, TC, and CHG.

## Statements and Declarations

### Ethical considerations

Protocols for animal experiments were approved by the Queen’s University Animal Care Committee (Protocol no. 2023-2457) on June 11^th^, 2024, in compliance with the Canadian Council on Animal Care (CCAC) guidelines for the care and use of laboratory animals.

No human participants were involved in this study.

### Data availability

The data that support the findings of this study are available from the corresponding authors upon reasonable request.

