## Supplemental Data for "Differential effects of BCG-Russia and BCG-TICE on trained immunity: potential implications for bladder cancer immunotherapy"

**Supplemental Table 1.** Raw values of plasma cytokine concentrations.

| IL-1β (pg/mL) | | |
|---|---|---|
| *PBS (n=5)* | *BCG-Russia (n=6)* | *BCG-TICE (n=4)* |
| 254.61 | 1535.99 | 5555.06 |
| 297.91 | 3777.92 | 3113.35 |
| 104.42 | 2617.33 | 2972.67 |
| 250.7 | 3610.47 | 1925.99 |
| 127.73 | 798.03 | |
| | 1440.93 | |
| **IL-12 (pg/mL)** | | |
| *PBS (n=5)* | *BCG-Russia (n=6)* | *BCG-TICE (n=4)* |
| 43.63 | 664.71 | 1300.73 |
| 142.96 | 1165.66 | 545.7 |
| 171.64 | 1314.05 | 650.92 |
| 142.96 | 1419.53 | 234.28 |
| 82.16 | 637.7 | |
| | 550.21 | |
| **TNF-α (pg/mL)** | | |
| *PBS (n=4)* | *BCG-Russia (n=5)* | *BCG-TICE (n=5)* |
| 53.62 | 1224.53 | 10032.08 |
| 97.62 | 1353.58 | 8797.36 |
| 109.70 | 2998.11 | 5536.23 |
| 50.83 | 1524.91 | 805.66 |
| | 537.74 | |
| **IL-10 (pg/mL)** | | |
| *PBS (n=5)* | *BCG-Russia (n=6)* | *BCG-TICE (n=4)* |
| 323.88 | 265.55 | 1139.5 |
| 506.06 | 584.63 | 251.25 |
| 78.21 | 84.96 | 696.9 |
| 66.03 | 325.18 | 427.51 |
| 549.75 | 276.82 | |
| | 361.34 | |
| **IL-2 (pg/mL)** | | |
| *PBS (n=4)* | *BCG-Russia (n=6)* | *BCG-TICE (n=4)* |
| 22.17 | 11.09 | 22.17 |
| 41.13 | 44.07 | 11.09 |
| 1.61 | 22.17 | 134.12 |
| 12.69 | 116.3 | 46.98 |
| | 39.01 | |
| | 60.71 | |
